# Defining the limits of plant chemical space: challenges and estimations

**DOI:** 10.1101/2025.01.08.631938

**Authors:** Chloe Engler Hart, Yojana Gadiya, Tobias Kind, Christoph A. Krettler, Matthew Gaetz, Biswapriya B. Misra, David Healey, August Allen, Viswa Colluru, Daniel Domingo-Fernández

## Abstract

The plant kingdom, encompassing nearly 400,000 known species, produces an immense diversity of metabolites, including primary compounds essential for survival and secondary metabolites specialized for ecological interactions. These metabolites constitute a vast and complex phytochemical space with significant potential applications in medicine, agriculture, and biotechnology. However, much of this chemical diversity remains unexplored, as only a fraction of plant species have been studied comprehensively. In this work, we estimate the size of the plant chemical space by leveraging large-scale metabolomics and literature datasets. We begin by examining the known chemical space, which, while containing at most several hundred thousand unique compounds, remains sparsely covered. Using data from over 1,000 plant species, we apply various mass spectrometry-based approaches—a formula prediction model, a de novo prediction model, a combination of library search and *de novo* prediction, and MS2 clustering—to estimate the number of unique structures. Our methods suggest that the number of unique compounds in the metabolomics dataset alone may already surpass existing estimates of plant chemical diversity. Finally, we project these findings across the entire plant kingdom, conservatively estimating that the total plant chemical space likely spans millions, if not more, with the vast majority still unexplored.

## 1. Introduction

Hundreds of thousands of species exist in the plant kingdom. Current estimates cover approximately 390,000 species, with a few thousand novel vascular plants being discovered every year (Enquist *et al*., 2019; KEW, 2016). Each plant produces thousands of primary and specialized metabolites for survival and environmental interaction. Thus, this vast and diverse phytochemical space can theoretically comprise millions of potential metabolites, some of which have a variety of applications (Kawatra *et al*., 2023). About 2% of these known plants have already been used for medicinal purposes (Domingo-Fernández *et al*., 2023).

Several studies have attempted to estimate the size of the chemical space for the plant kingdom, emphasizing the complexity of plant metabolism. These metabolites can be classified into two global categories: the primary core metabolites that are broadly shared across all species, and secondary metabolites, which are specialized plant compounds. Around 8,000 metabolites that reoccur across multiple species are captured in databases such as PlantCyc (Hawkins *et al*., 2024). Secondary plant metabolites that have been identified from the literature have been captured in open natural product (NP) databases such as COCONUT (Chandrasekhar *et al*.,2024) and LOTUS (Rutz *et al*., 2022), which cover approximately 125,000 plant-based compounds. These databases provide a foundation for understanding the diversity of plant metabolites, though they represent only a fraction of the vast chemical space yet to be explored.

Early plant metabolomics research estimated that there are 200,000 plant-derived compounds based on the hypothesis that each species produces at least five novel secondary metabolites and the assumption that approximately 223,000 plant species were known at the time (Afendi *et al*., 2011; Scotland and Wortley, 2003). It is worth noting that the plant chemical space is a subset of the entire theoretical chemical space, which is much larger and is further reviewed for the interested reader in **Supplementary Text 1**.

The identification of chemical structures in complex samples, such as plant extracts, has improved with advancements in metabolomics. However, mapping the entire phytochemical space remains a difficult task. Untargeted metabolomics methods, including liquid chromatography/mass spectrometry (LC/MS), still rely on deep sampling across various enrichment and separation techniques. The quickest method for compound annotations is mass spectral library search (Bittremieux *et al*., 2022). Nevertheless, such mass spectral libraries are hampered by the small number of reference mass spectra, specifically in the NP space. A wide array of computational mass spectrometry tools have been developed to support the structure elucidation process (Krettler and Thallinger, 2021), including algorithms that utilize fingerprint lookups in databases of known compounds such as CSI:FingerID (Dührkop *et al.,* 2015). The most promising method for identifying metabolites that can not be found in any database yet is *de novo* machine learning algorithms such as MS2Mol (Butler *et al*., 2022). Nonetheless, structure verification for many novel compounds still requires the help of nuclear magnetic resonance (NMR). Unfortunately, NMR can not easily be scaled to achieve a similar throughput to LC/MS due to the low sensitivity and very low sample throughput.

In this work, we leverage one of the largest publicly available metabolomics and literature datasets to estimate the chemical space of the plant kingdom. We begin by examining the known chemical space and find that only a few tens of thousands of plant species have been studied, most of them only superficially. The chemical space documented in the literature likely contains several hundred thousand unique structures, providing only a glimpse of the true diversity. We then analyze the overlap between the metabolomics and literature datasets for plants present in both, observing only a moderate alignment. This suggests that a single or a few metabolomics samples per plant cannot capture the full metabolome. Subsequently, we predict the number of unique chemical structures in a metabolomics dataset spanning over 1,000 plant species using various complementary approaches: i) a formula prediction model, ii) a *de novo* prediction model, iii) library-based search combined with the *de novo* model, and iv) MS2 clustering. Our results indicate that the number of unique structures in this dataset may already exceed current estimates of the phytochemical space. Finally, we project these findings to estimate the total size of the chemical space across the entire plant kingdom, suggesting that it likely spans into the millions. These projections more than likely indicate that over 99% of the phytochemical space remains unexplored, highlighting its vast and largely untapped potential.

## 2. Methods

### 2.1. Collecting publicly available mass spectrometry data from plants

We identified several of the largest datasets publicly available through the Experimental Natural Products Knowledge Graph (ENPKG), a Knowledge Graph structured in a Resource Description Framework (RDF) to harmonize heterogeneous NP metabolomic datasets (Gaudry *et al.,* 2024). Apart from harmonizing several datasets and allowing query them, ENPKG contains thousands of associations between LC/MS features and their corresponding metadata, such as their SIRIUS/CSI:FingerID-predicted structures (Dührkop *et al.,* 2015; Dührkop *et al.,* 2019), references to the ISDB-LOTUS database (https://zenodo.org/records/7534250), and references to their derived extracts. These metadata are externally linked using standard identifiers such as WikiData for extract species, chemical class through NPClassifier (Kim *et al*., 2021), and predicted structure through SMILES, InChIKeys, and ChEMBL identifiers (Gaulton *et al*., 2012). Throughout the manuscript, we refer to chemical structures as plant metabolites interchangeably. The current version of ENPKG integrates three medicinal plant datasets and three bacterial datasets (*L. donovani, T. cruzi,* and *T. brucei rhodesiense).* We restricted our analysis to the three medicinal plant datasets **(see details in Table 1**).

**Table 1.** Overview of the three medicinal plant datasets in ENPKG. The first column references the MassIVE ID of the dataset, where the metabolite spectra can be found. The second and third columns are the plant library and its original publication. The last three columns report the number of extracts in the library, the number of unique structures (represented by SMILES) predicted by CSI:FingerID, and the number of features detected. Note that the number of extracts does not correspond to the number of plant species, as the datasets could have multiple extracts for one species.

| MassIVE ID | Plant Library | Reference | # Extracts | # Unique structures | # Features |
| --- | --- | --- | --- | --- | --- |
| MSV000093464 | Korean Pharmacopoeia Plants Extracts | Kang *et al.*, 2022 | 337 | 17,428 | 42,520 |
| MSV000088521 | *Waltheria Indica* Samples (aerial parts and roots) | Cretton *et al.*, 2016; Cretton *et al.*, 2014 | 1 | 2,453 | 4,422 |
| MSV000087728 | Pierre Fabre Research Institute Library (RFRIL) | Allard *et al.*, 2024 | 1,600 | 65,381 | 784,836 |

We queried the ENPKG (accessed on 26 November 2024) via their SPARQL endpoint in GraphDB (https://enpkg.commons-lab.org/graphdb/) to extract all LC/MS features with the following annotations: (a) an InChIKey, (b) a SMILES, (c) NP class, superclass, and pathway annotation from NPClassifier, (c) structural annotation from CSI:FingerID and confidence scores, (d) predicted formula from SIRUS, and (e) a WikiData species annotation. The SPARQL query used for this extraction is provided in **Supplementary Text 2**. This resulted in 76,982 unique chemical structures and 831,778 features across 1,018 unique plant species (Figure 1A). We would like to note that we used a consistent arbitrary ordering of these plants in our analyses, but we confirmed that the ordering does not significantly affect the results **(Supplementary** Figure 1). Finally, we downloaded the raw spectra for these features using MassIVE’s API (https://massive.ucsd.edu/).

**Figure 1.**
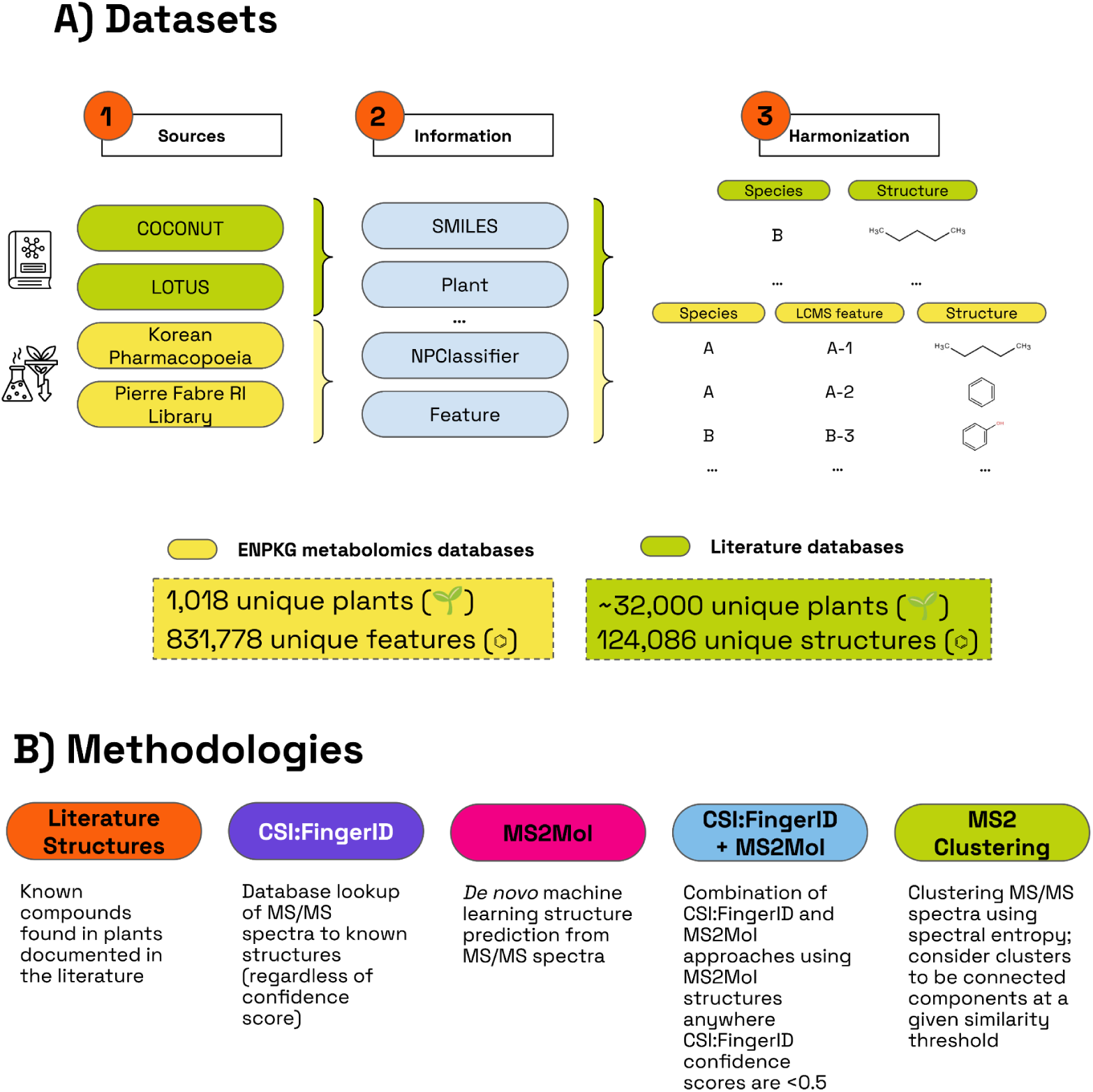
A) Datasets leveraged in our work and the corresponding information extracted from them. B) Methodologies employed to calculate the number of unique chemical structures in the datasets.

The 831,778 features in the dataset correspond to various adduct types **(Supplementary Table 1)**. To reduce potential redundancy, we filtered out all features not predicted as [M+H]+ adducts by SIRIUS (Dührkop *et al.,* 2019), resulting in 335,377 features. While this step excludes over 50% of the features and disregards negative mode samples, it significantly reduces the risk of overestimating the projections for the chemical space. Notably, we did not apply this filtering step to the formula prediction method since it accounts for adducts.

We further categorized the predicted structures into known and unknown groups based on an arbitrary CSI:FingerID confidence score threshold of 0.5. These confidence scores, generated using COSMIC (Hoffmann *et al.,* 2022), have been proven to separate correct and incorrect structures (using the CASMI 2016 dataset). About 75% of the features from the dataset were classified as unknown (34,808 chemical structures), while the remaining 25% were categorized as known (12,434 chemical structures). The rationale behind this classification was to annotate later unknown features (i.e., low confidence annotation), which are likely not in the reference library, using a *de novo* structured prediction model. Figure 2 details the confidence scores, each group’s features percentage, and MS precursor mass distribution. Supplementary Figures 2 and 3 also show the distribution of the predicted NP super class and classes using CANOPUS (Dührkop *et al*., 2021).

**Figure 2.**
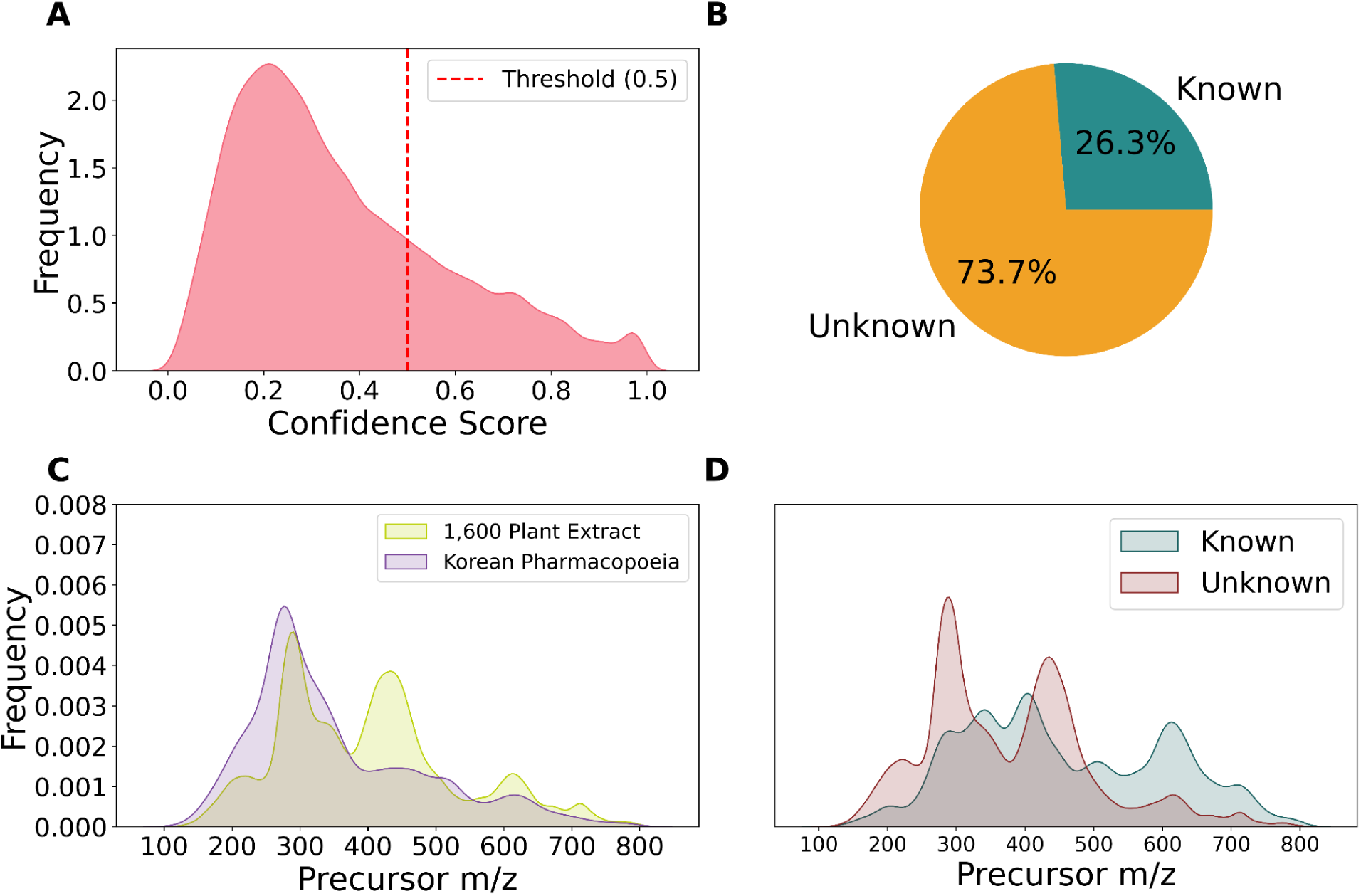
Summary of the ENPKG dataset. A) Distribution of the CSI:FingerID confidence scores for the predicted structures across all features with a predefined threshold of 0.5 (dotted red line) to classify confidently predicted (known) and unknown features. B) Distribution of confidently predicted (known) and unknown features in the dataset. C) Distribution of the precursor m/z across all the features stratified by the two primary datasets. D) Distribution of the precursor m/z for known and unknown features in the dataset.

Lastly, we note that the quality of the public datasets used constrains the methods described below for estimating the size of chemical space. Notably, we found that 1.04% of the spectra in the ENPKG dataset had a dominating peak with more than 90% of the total intensity. Such spectra are generally uninformative and may not have been accurately classified by predictive models or MS2 clustering methods. To help mitigate this issue, we removed any spectra from the dataset with less than five peaks before creating the data curves.

### 2.2. Harnessing chemical structures reported in plants across the scientific literature

While the mass spectrometry datasets described in the previous section provide a snapshot of the metabolites present in a specific plant under specific conditions, the LC/MS features identified for a given plant are far from representing all actual metabolites present in the sample. Without going deep into the topic, which is mentioned later in the discussion, various technical factors affect the detected chemical space of a given sample, such as chromatography column, extraction method, etc. Furthermore, different plant parts will produce distinct metabolites, and the abundance of a metabolite in the mixture can be, in some cases, too low to be detected. Thus, we also complement our analysis with two of the largest datasets comprising metabolites that have been reported in the natural world: COCONUT 2.0 (Chandrasekhar *et al*., 2024) and LOTUS (Rutz *et al*., 2022) (Figure 1A).

While both datasets comprise multiple sources of information, given our goal, we simplified the datasets to the associations between chemical structures and their reported taxonomic species. The datasets capture this information in two columns: SMILES and species name.

However, given that both datasets contain other taxonomic clades apart from plants, we assigned all taxonomic species to a unique identifier of the following plant taxonomic nomenclatures: NCBITaxonomy (Cox *et al*., 2024), Integrated Taxonomic Information System (ITIS) and World Flora Online (WFO). For NCBITaxonomy, we subset exclusively to all species under the Viridiplantae kingdom (NCBITaxon:33090).

To avoid the indistinctive assignment of species to different nomenclatures, we established a prioritization order, namely: (i) NCBITaxon, (ii) ITIS, and (iii) WFO. Thus, if a species name is present in several nomenclatures, it is exclusively assigned to the identifier of the nomenclature with the highest priority. After applying this harmonization approach, this prioritization is reflected in the final distribution of plants captured in both datasets **(Supplementary Table 2)**. Lastly, we parsed SMILES with RDKit (Landrum, 2016) and converted them to InChIKeys (first 14 characters) while filtering out invalid SMILES. The number of unique structures identified was 124,086.

### 2.3. Predicting formulas and structures from MS2 data

We used three methods to calculate the unique number of structures and generate structure curves from MS2 data (Figure 1B). Firstly, we also employed a *de novo* model, MS2Mol (Butler *et al*., 2022), which can predict novel structures in previously unexplored regions of chemical space. Secondly, we used a hybrid approach combining the confident predictions from CSI:FingerID (i.e., known group) with the MS2Mol predictions for the remaining spectra. Thirdly, we predicted the formulas (which can lead to multiple structures) for each feature to establish a lower bound using SIRIUS, a state-of-the-art formula prediction model (Dührkop *et al.,* 2019), acknowledging that each formula corresponds to multiple possible structures.

### 2.4. Spectral clustering

As an alternative method for estimating the number of structures, we implemented MS2 clustering to avoid dependence on predictive models (Figure 1B). Using the spectral entropy metric (Li *et al*., 2021), we constructed a similarity matrix for the spectra in our dataset. We limited comparisons to spectra with precursor m/z within 10 and 200 ppm to minimize false matches, assigning a similarity score of zero to all other spectral pairs. While the results in this paper are based on a 10 ppm filter, no significant changes were observed when using a 200 ppm filter. Clustering was performed by thresholding the similarity scores at 0.5, 0.6, 0.7, 0.8, and 0.9 (**Supplementary** Figure 4). The paper primarily relies on results using thresholds of 0.7 and 0.8, aligning closely with the 0.75 threshold reported by Li *et al*. (2021) to have a false discovery rate of less than 10%. Finally, we constructed a network with *igraph* (Antonov *et al*., 2023) and identified clusters based on the connected components.

### 2.5. Projecting the total chemical space

Applying the previously described estimation methods to the public ENPKG dataset gave us estimates for the number of plant metabolites in 1,000 plants. To project the total number of plant metabolites across all 400,000 plant species, we fitted power law models (**Supplementary Function 1**) to the data. We employed holdout sets of 30%, 20%, and 10% to validate our predicted curves, observing minor variations (**Supplementary** Figure 10).

### 2.6. Implementation

All scripts used in this work were written in Python version 3.10 and managed using the Poetry environment (https://python-poetry.org/). For data manipulation, we employed widely used libraries such as Pandas (McKinney, 2010), NumPy (Harris *et al.,* 2020), and SciPy (Virtanen *et al.,* 2020). Visualizations were generated using seaborn (Waskom, 2021) and Matplotlib (Hunter, 2007). To represent chemical structures, we first employed SMILES that were converted to InChIKeys using RDKit (Landrum, 2016). Notably, we represented chemical structures using the first block of 14 characters from the InChIKey (out of a total of 27) to intentionally exclude stereochemistry, since considering it would significantly inflate estimations. For MS2 data processing, we relied on specialized libraries, including MatchMS (De Jonge *et al*., 2004). The clustering of MS2 data was performed using the *igraph* library (Antonov, 2023). Lastly, this work’s data and source code are publicly available on Zenodo and GitHub at https://github.com/enveda/chemical-space-estimation.

## 3. Results

### 3.1. Exploring the known chemical space of the plant kingdom

Determining the exact size of the chemical space of the plant kingdom remains unattainable with current knowledge. However, we can make the most accurate estimate possible using publicly available data. To achieve this, we first evaluated the known chemical space of the plant kingdom by constructing a chemical space saturation curve using the two largest publicly available literature datasets (Rutz *et al*., 2022; Chandrasekhar *et al*., 2024) (Figure 3A). As expected, the cumulative curve shows a decreasing growth rate as more plants are added, eventually plateauing at approximately 124,000 unique structures for the 32,000 species with available data. Additionally, we examined this trend using Murcko scaffolds and observed a similar pattern **(Supplementary** Figure 5). Interestingly, both curves exhibit sharp increases in certain regions, reflecting instances where the addition of a species contributed a disproportionately large number of new structures. This phenomenon highlights biases in the literature, where a few extensively studied species account for thousands of reported structures, whereas most species have only a few, if any, structures documented.

**Figure 3.**
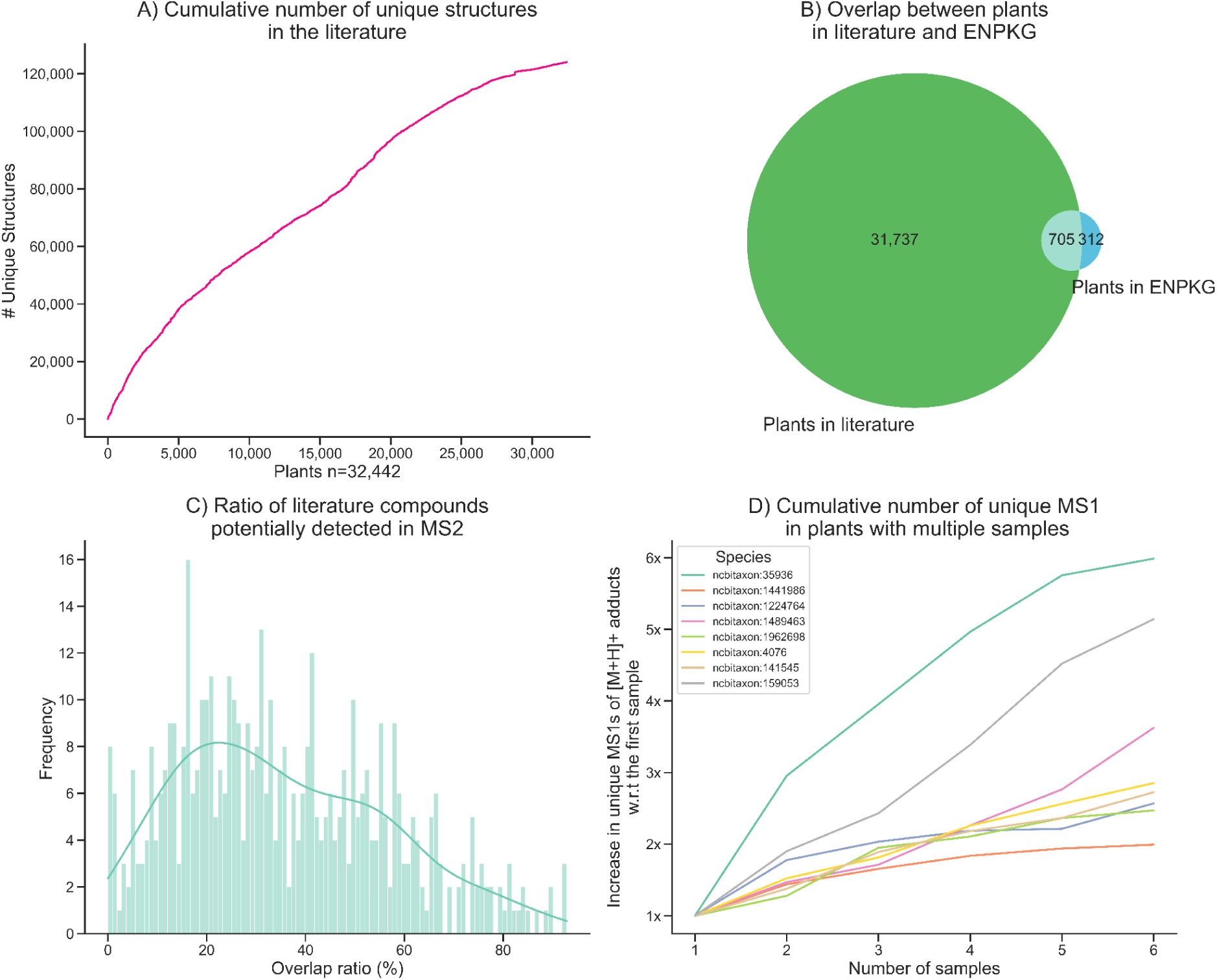
A) Cumulative curve of unique chemical structures in the literature dataset. B) Overlap between the plants in the literature and the ENPKG datasets. B) Overlap between plants in literature and ENPKG. C) Distribution of the ratio of literature compounds that can be potentially detected in MS2 spectra based on all the precursor mz mass shifts. Supplementary Table 3 lists the adducts used to calculate each feature’s potential precursor mz mass shifts. **D) Cumulative number of unique precursor masses (MS1) in plants with multiple samples.** The Y axis shows the X increase in MS1s as more samples are added. The eight plants used are the only ones containing more than five samples in ENPKG.

Before estimating the plant chemical space using mass spectrometry data, we first evaluated the coverage of the mass spectrometry dataset in relation to the literature dataset, given their substantial overlap of plants (Figure 3B). Unfortunately, the literature dataset is sparse, with most plants represented by only a few dozen unique structures **(Supplementary** Figure 6). Therefore, we focused our analysis on 490 plants present in both datasets with 20 or more unique structures. As a proxy to assess coverage, we calculated all potential precursor m/z mass shifts based on the adducts listed in **Supplementary Table 3** and compared them to the exact masses of compounds reported in the literature for the same plant (Figure 3C). It is important to note that the MS2 dataset does not capture the entirety of the metabolites present in the plants, further constraining the observed overlap. The overlap in Figure 3B represents the percentage of chemical structures in the literature for which a matching precursor m/z mass shift exists, indicating that these compounds could be identified in the metabolomics dataset. Since MS2 signals could correspond to other structures, the observed overlap represents an upper limit of the coverage, as some matches may not directly correspond to the exact compounds reported in the literature. Conversely, the absence of a precursor m/z mass shift matching a given compound indicates that the compound is not present in the metabolomics dataset. The overlap ratios across these 490 plants follow a normal distribution centered around 25%, indicating that for most plants, the metabolomics dataset can potentially capture only a moderate proportion of the compounds reported in the literature (Figure 3C). This confirms that a substantial number of metabolites documented in the literature are not present in the metabolomics dataset.

Another aspect we were interested in investigating is how the coverage of the metabolome increases as more samples are screened for a given species. To do so, we calculated the cumulative number of unique precursor masses (MS1s) rounded to two decimals for eight species containing more than five samples in ENPKG (Figure 3D). The results show how going from a single sample to six increases the metabolome coverage up to six times. These findings suggest that while the ENPKG dataset includes over 1,000 plant species, it may capture only a portion of their true chemical diversity. As a result, the estimates derived from this dataset might not entirely reflect the metabolomic richness of these plants, given the current sampling depth.

We also assessed the consistency of metabolite coverage for extracts from 15 plants shared between the Korean Pharmacopoeia (Kang *et al*., 2022) and the Pierre Fabre Research Institute Library (RFRIL) (Allard *et al.,* 2024), the two main extract libraries within ENPKG. To do this, we compared the distribution of precursor masses and the overlap of predicted Murcko scaffolds from CSI:FingerID across both libraries **(Supplementary** Figure 7). Our analysis revealed noticeable differences in precursor masses and a low overlap in predicted scaffolds between the two libraries. Additionally, the RFRIL samples contained a larger number of features compared to the extracts from the Korean Pharmacopoeia. These discrepancies in precursor masses and scaffold overlap likely stem from differences in extraction methods and other factors influencing the metabolites captured in a metabolomics dataset, which are further examined in the discussion section. Overall, the multiple findings here suggest that the estimates presented in the following subsections likely underestimate the true size of the plant metabolite space.

### 3.2. Public mass spectrometry data may already surpass current estimates of phytochemical space size

To assess how many unique chemical structures may already exist in current datasets, we leveraged one of the largest publicly available metabolomics datasets for plants (ENPKG) (Gaudry *et al.,* 2024). We used a combination of four approaches to get these estimates: i) predicted formulas (SIRIUS), ii) de novo modeling (MS2Mol), iii) hybrid (CSI:FingerID + MS2Mol), and iv) MS2 clustering. We selected MS2Mol since it was one of the first publicly available de novo models, and we developed it. We also selected CSI:FingerID because it serves as the standard library reference model. Alternatively, other de novo models such as MSNovelist (Stravs *et al*., 2022) or database lookup approaches (e.g., spectral entropy or cosine similarity) could have been used. These choices allowed us to get varying estimates from three widely used approaches with different strengths and limitations.

Our results reveal that the *de novo* model predicts the most structures, surpassing 100,000 unique structures across the 1,000 plants. In contrast, the number of unique formulas is the lowest due to its inherent constraint (Figure 4). MS2 clustering and the hybrid approach (CSI:FingerID + MS2Mol) yield estimates that fall between these two extremes. Interestingly, while the formula curve is very likely a substantial underestimate since one formula can correspond to multiple structures, it is higher than the MS2 clustering curve at the beginning, suggesting that these curves may also be conservative. These findings highlight the differing capabilities and scopes of the models. Rather than directly comparing the methods — which is not the focus of this work — we emphasize that the predictions from the de novo model, the hybrid approach, and MS2 clustering are close to the size of the literature (124,000 metabolites) with far fewer plants. Overall, given that there is mass spectrometry data for 3,000 plants in GNPS and other public repositories and the curves increase linearly, it is more than likely that current mass spectrometry data may exceed current phytochemical space size estimates.

**Figure 4.**
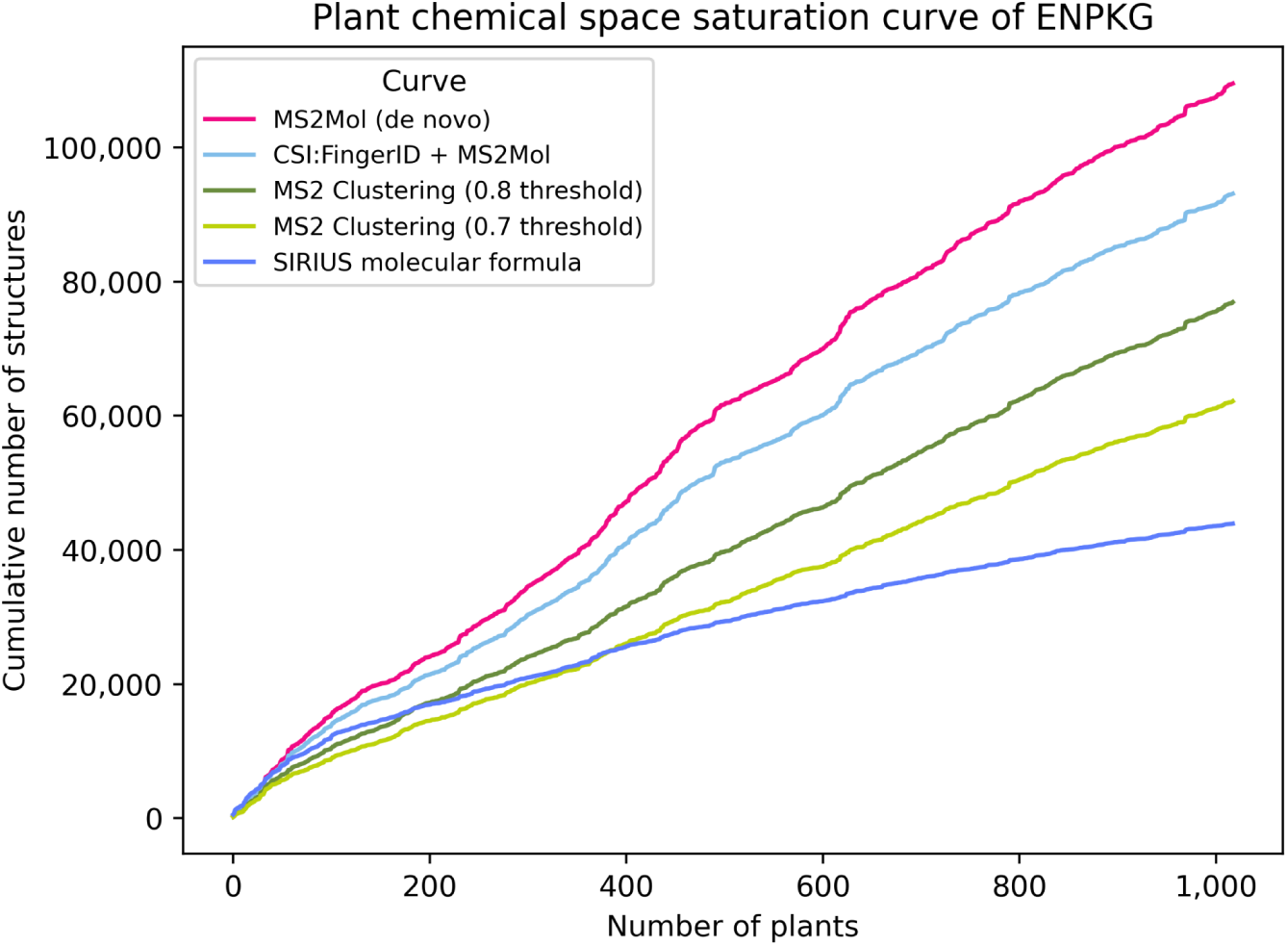
Plant chemical space saturation curve of ENPKG. The plot shows a cumulative curve of the predicted number of unique structures using different methodologies. These curves are used later to fit power law models to estimate the total phytochemical space.

Despite these differences in cumulative estimates, the structure curves are strikingly similar in shape, with steeper increases at consistent points across methods. This indicates that certain plants consistently contribute new chemicals regardless of the approach used. We note that the taxonomic diversity in this public dataset is high **(Supplementary Table 4)**, which likely contributes to the curves’ steepness. Thus, we believe a dataset with 1,000 less diverse plants might have shown slower increases.

Additionally, we observed similar trends when analyzing Murcko scaffolds instead of structures **(Supplementary** Figure 8). These patterns suggest the models are not simply inflating the numbers based on specific regions of the chemical space but are capturing trends across different plants. To further validate this, we compared the predicted structures for a given MS2 spectrum between the two models **(Supplementary** Figure 9). We observed a relatively high level of agreement, supporting the consistency and reliability of the predictions.

### 3.3. Projections of the plant chemical space for plants range from millions to tens of millions

To estimate the total number of plant metabolites, we fit power law models to the curves in Figure 4 and projected them to 400,000 plants, yielding estimates between 1.5 and 25.7 millions total plant metabolites (Figure 5). The formula curve probably underestimates the size, as a given formula can lead to multiple structures. On the other hand, the MS2Mol curve is affected by the accuracy of the predictions from the model, which was not necessarily trained to get exact matches. Similarly, the hybrid model resembles the MS2Mol curve since most CSI:FingerID predictions have low confidence (Figure 2A). Given these limitations of the predictive models, the MS2 clustering methods are, in our opinion, the most appropriate for this task.

**Figure 5.**
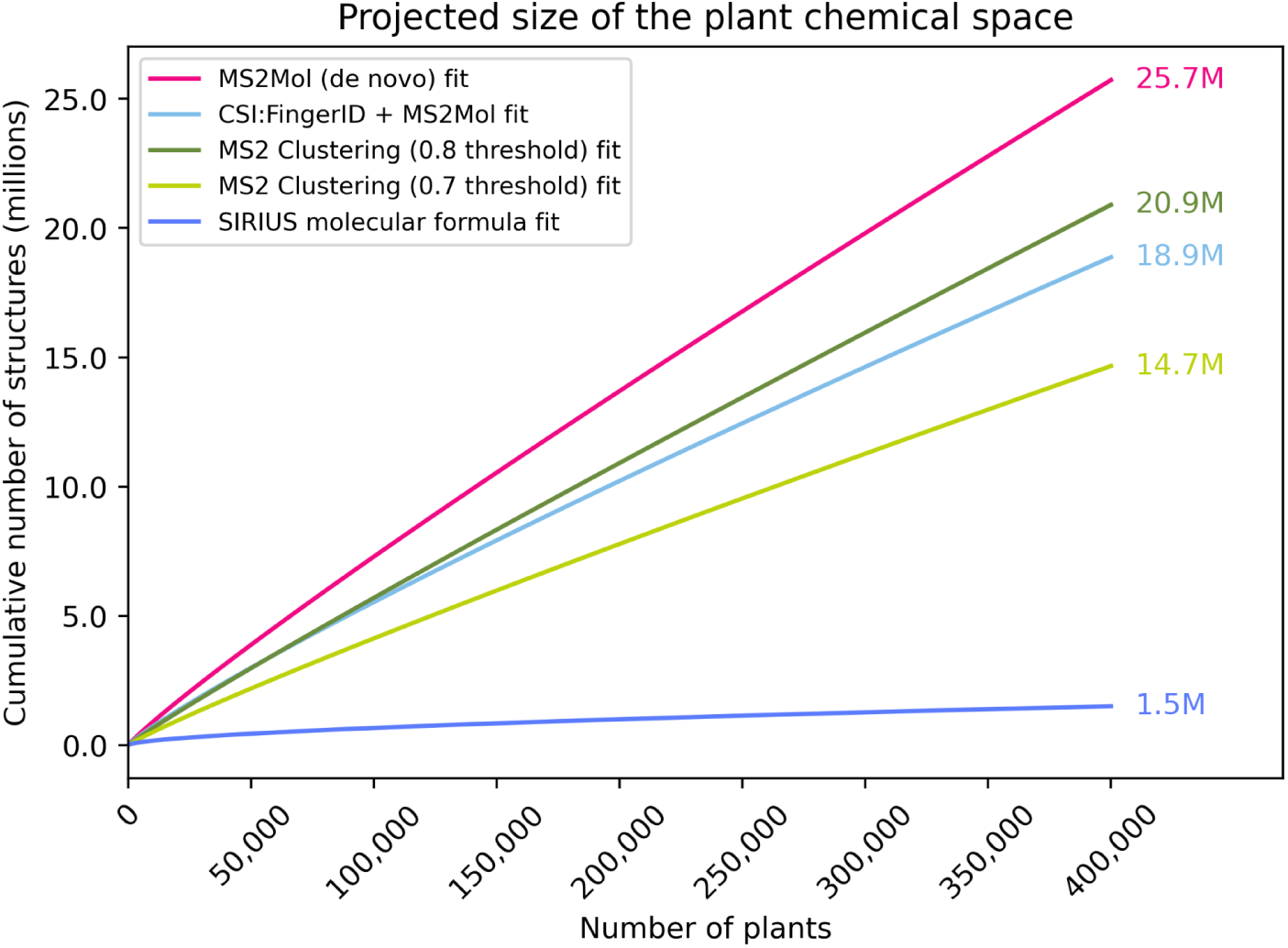
Estimated size of the chemical space after fitting the power law curves across various methodologies. The numbers on the right side correspond to the size of the chemical space in millions for each method, assuming there are 400,000 species of plants. **Supplementary** Figure 11 depicts a zoomed-in view of the plot around the first 1,000 species.

Additionally, we estimated the number of metabolites for 35,000 plant species, a value comparable to the number of species documented in the literature, to evaluate how our projections align with currently available data. For the MS2Mol model, the predicted number of unique structures approaches 3 million, while MS2 clustering yields estimates ranging from 1.6 to 2.1 million, depending on the similarity thresholds of 0.7 and 0.8, respectively. These findings highlight the significant gap between the predicted metabolite diversity and the literature’s current coverage, emphasizing how much of the plant chemical space remains unexplored.

Lastly, we used holdout sets comprising the last 10%, 20%, and 30% of the data to validate our models. These sets resulted in minor variations in how well the curves fit the data **(Supplementary Figure 10A)**, magnified when the curves were projected for 400,000 plants **(Supplementary Figure 10B)**. These variations were most pronounced for the CSI:FingerID curves since they flattened out more quickly than the others. This suggests that as more data becomes publicly available, these data curves may change and plateau sharply.

In conclusion, our estimates reveal that the lower bound for potential plant metabolites is higher than previously estimated, exceeding earlier estimates by a factor of at least 10. However, the current datasets constrain the upper and lower bounds and could evolve as more public data becomes available.

## 4. Discussion

Our work relies on multiple factors that can significantly influence the resulting estimates; therefore, discussing the most critical considerations is crucial. **Table 2** summarizes the limitations discussed in this section.

**Table 2.** Summary of the major limitations of our work.

| Limitation | Summary | Effect on Estimates |
| --- | --- | --- |
| **Dataset coverage** | Estimates are based on a small fraction of plant species, leading to potential biases | Unknown |
| **Extraction and chromatographic bias** | Different protocols and incomplete anatomical plant coverage may omit specific metabolites | Underestimation |
| **MS2 Spectrum prediction accuracy** | Library constraints and clustering methods may | Unknown |
|  | overestimate or fail to generalize |  |
| **Disparate solvents used** | Different solvents extract different compounds, causing inconsistencies | Underestimation |
| **Adducts considered** | Only [M+H]<sup>+</sup> adduct was considered | Underestimation |
| **Missed compound classes** | Polar and moderately polar methods miss volatile and lipid compounds | Underestimation |
| **False positives in dataset** | Annotation errors and contaminants may inflate metabolite counts | Overestimation |
| **Functional genomics potential** | Genomics tools could dramatically expand estimations but remain underexplored | Underestimation |
| **Projection modelling** | Power law projections may overestimate space if actual data curves plateau | Overestimation |

First, although we employed one of the largest publicly available datasets, our estimations are based on a metabolomics dataset that represents less than 1% of all species in the plant kingdom. While including the literature dataset expanded the coverage to approximately 10%, only a small fraction of those plants have been studied comprehensively. This limited coverage introduces potential biases, as the plants that have been extensively studied are not necessarily representative of the vast chemical diversity present across the entire plant kingdom. Consequently, our estimates are influenced by the depth and breadth of current data, highlighting the need for further profiling and exploration to better capture the true extent of plant chemical diversity. These limitations are particularly pronounced when projecting curves using power law models to get estimates for 400,000 plants, as the actual data curves may plateau beyond 1,000 plants, leading to lower estimates.

Second, our estimations are derived from metabolomics data obtained through specific extraction methods and chromatographic protocols, which may not capture a substantial portion of the metabolome (discussed later in detail). For instance, the ENPKG data was generated using different extraction techniques and chromatography columns, such as the BEC C18 non-polar column. Employing alternative protocols could uncover undetected metabolites under the current experimental settings. Additionally, our analysis lacks comprehensive coverage of plant extracts from all anatomical parts—such as roots, stems, and leaves—for most species. This limitation is critical, as metabolite production varies significantly between plant parts and is further influenced by environmental conditions (e.g., season, stress, temperature, humidity, etc.) (Figure 3D). These factors collectively constrain the comprehensiveness of our estimations and suggest that the true diversity of plant metabolites may be even greater than what our analyses indicate (Figure 3C-D).

For instance, Aghdam and Brown (2021) estimated that the size of the chemical space of endophyte-derived secondary metabolites could be in the range of several billion unique compounds. These estimates were derived using assumptions such as ∼90% of secondary metabolite biosynthetic capacity being silent or cryptic, the presence of approximately 10 host-specific bacterial endophytes per plant species, and hundreds of fungal endophyte species per plant. Their approach also relied on rough approximations of species-to-metabolite ratios. While these estimates are impressive, the assumptions may be optimistic since every endophyte species is unlikely to produce entirely distinct metabolites, as overlap and redundancy in biosynthetic pathways are common. Additionally, their calculations assume that each endophyte species acts independently, disregarding interactions between microbes, plants, and environmental factors that may influence metabolite production.

The third major factor is the accuracy of the predicted structures for each MS2 spectrum. Library-based models like CSI:FingerID rely on a fixed library (∼1 million biomolecules), while generative models trained on mass spectrometry data struggle to generalize beyond the chemical space in their training (Engler Hart *et al*., 2024). To address these limitations and focus solely on the number of unique structures in the dataset, we clustered the MS2 spectra to identify groups corresponding to the same chemical structure. This approach, however, is sensitive to the spectral entropy threshold chosen to define the clusters **(Supplementary** Figure 4) and may cluster spectra from distinct molecules with similar patterns, especially if they have few peaks. Additionally, differences in the MS/MS acquisition methods– Kang *et al*. (2022) used a collision energy ramp from 20-100 eV, while Allard *et al*. (2024) used three distinct collision voltage settings (15, 30, and 45eV)--may lead to overcounting unique structures because they may be represented by slightly different MS/MS fragmentation spectra. To mitigate these issues and avoid overestimation, we made our estimations exclusively relying on the predominant adduct ([M+H]+) and filtered out spectra with less than five peaks. However, this likely underestimates the true chemical diversity, as other adduct forms, which may represent unique metabolites, are excluded from consideration.

The fourth factor is the disparate solvents used for plant extraction (Vijayan and Chandra, 2015). The Korean Pharmacopoeia dataset (Kang *et al*., 2022) used a methanolic extract. Methanol is a polar solvent and extracts a wide range of compound classes (alkaloids, flavonoids, phenolic acids, and tannins). The second dataset (Allard *et al.,* 2024) utilized ethyl acetate as the extraction solvent, a moderately polar solvent leading to the extraction of sterols, terpenoids, and less tannins. The different solvents used can explain the differences observed among the 15 extracts of plants in both datasets **(Supplementary** Figure 7). The fifth factor is that the extraction methods above will not capture many other compound classes and require different measurement technologies. The polar and moderately polar extraction and the LC-MS-based method ignore numerous other substance classes, such as volatile compounds and lipids. For Volatile Organic Compounds (VOCs) gas-chromatography coupled to mass spectrometry (GC-MS) has to be used. For polar and neutral lipids, specialized lipidomic extraction techniques have to be applied.

The sixth factor relates to false positive compounds in the dataset, such as annotation errors from CSI:FingerID and MS2Mol. For example, plasticizers or other contaminants such as pesticides. While they can be considered correct for sample annotation, they can not be considered plant metabolites (Kind *et al*. 2015). For example, we have observed several thousand fluorinated and chlorinated SMILES structures in annotations from both models. To confirm the presence or absence of such compounds, one would need to run additional analytical tests, which go beyond the scope of this paper. The seventh factor is a further path that relies on functional genomics approaches by massively increasing the compound space for compounds of interest. That can be done by CRISPR/Cas-based gene editing and other advanced techniques, including cytochrome P450 transform in plants (Nguyen *et al*. 2021).

In summary, while our study provides a foundational estimate of the size of the plant chemical space, significant opportunities remain for future research to refine and improve these projections. Notably, most factors listed in **Table 2** suggest our estimates are likely conservative and lean towards underestimation. These limitations highlight the need for broader datasets and experimental approaches to capture more of the plant metabolome space. As more data becomes available through sources like MetaboLights (Haug *et al*., 2020), Metabolomics Workbench (Sud *et al*., 2016), and Global Natural Products Social Networking (GNPS) (Wang *et al*., 2016), the curves can be regenerated and the estimates adjusted accordingly.

## 5. Conclusion

In this study, we sought to estimate the size of the chemical space of plants by leveraging one of the largest publicly available metabolomics and literature datasets. While deriving an exact number remains infeasible, we employed multiple approaches to establish a robust estimate of the potential range of the plant chemical space under various assumptions. First, we examined the known chemical space by analyzing the saturation curve derived from 124,000 unique chemical structures documented across over 30,000 plant species in the literature. Our findings highlight that the chemical space exhibits signs of saturation as additional species are studied, reflecting most plants’ limited exploration depth.

Next, we extended our analysis to metabolomics datasets comprising over 1,000 plant species. By leveraging mass spectrometry data, we provided a broader perspective on how the chemical space expands with increased profiling. Notably, the estimates for the number of unique structures across the 1,000 plants analyzed are already close to previous estimates of the chemical space (Scotland and Wortley, 2003; Afendi *et al*., 2011) and the number of structures documented in the literature. Furthermore, after we modeled growth curves to extrapolate the size of the chemical space based on an estimated 400,000 plant species, even the most conservative estimates reached into the millions. Based on molecular formulas, we estimated 1.5 million unique compounds, which, while relatively reliable, almost surely underestimate the true diversity. Additionally, the different structure prediction and MS2 clustering approaches estimated at least 15 million unique structures. Collectively, these findings suggest that plants’ chemical space could range from several million to tens of millions of unique structures.

## Authors’ contributions

CEH and DDF designed the study. CEH, YG, and DDF prepared the datasets. CAK generated the formula prediction dataset. CEH and DDF analyzed the datasets. CEH, YG, and DDF made the figures. CEH, TK, and DDF interpreted the results. CEH, YG, TK, and DDF wrote the paper. All authors reviewed the manuscript. All authors have read and approved the final manuscript.

## Funding

No funding is applicable.

## Acknowledgments

We are grateful to the authors of the ENPKG work, particularly Pierre-Marie Allard and Arnaud Gaudry, for their valuable assistance in answering our questions.

## Competing interests

All authors were employees of Enveda during the course of this work and have real or potential ownership interest in the company.

## Key points

- The plant kingdom’s chemical diversity is immense, with millions of unique metabolites estimated, yet remains largely unexplored.
- Mass spectrometry-based analyses reveal that known plant chemical diversity significantly underestimates the true scale of unique compounds.
- Our projected estimates suggest that the plant chemical space spans millions of metabolites, highlighting untapped potential for applications in medicine, agriculture, and biotechnology.

