## Supplementary File for "Defining the limits of plant chemical space: challenges and estimations"

#### Supplementary Text

**Supplementary Text 1. Literature review on estimating the entire size of the chemical space**

Several studies have attempted to estimate the size of chemical space, each employing different considerations and constraints to define its boundaries. Reymond *et al.* (2015) referenced an estimate of $10^{60}$ for "drug-like" chemical space, defined as molecules obeying Lipinski's rule of five for oral bioavailability. This estimate restricts elements to C, H, O, N, and S, assumes a maximum of 30 atoms to stay below 500 daltons, allows for branching, and permits a maximum of 4 rings. Polishchuk *et al.* (2013) estimated the number of realistic drug-like molecules that could potentially be synthesized to be around $10^{33}$, a more extensive estimation compared to Ertl (2003) ($10^{23}$). Drew *et al.* (2011) took a more conservative approach, estimating the size of organic chemical space to be approximately $3.4 \times 10^9$ molecules while also predicting the known drug space to be around $2.0 \times 10^6$. In conclusion, these estimates highlight the complexity of defining chemical space and demonstrate how different constraints and methodologies can lead to vastly different results, ranging from $10^9$ to $10^{60}$ molecules depending on the specific parameters.

**Supplementary Text 2. SPARQL Query to extract data from ENPKG.**

```
Unset
PREFIX enpkg: <https://enpkg.commons-lab.org/kg/>
PREFIX rdf: <http://www.w3.org/1999/02/22-rdf-syntax-ns#>

SELECT DISTINCT ?ik_2d ?smiles ?adduct ?np_class ?np_superclass
?np_pathway
?cosmic ?species ?feature ?feature_list ?massive_id ?row_id
?sample ?mode
WHERE
     {
          ?ik_2d rdf:type enpkg:InChIkey2D .
          ?ik_2d enpkg:has_smiles ?smiles .
          ?ik_2d enpkg:has_npc_class ?np_class .
          ?ik_2d enpkg:has_npc_superclass ?np_superclass .
          ?ik_2d enpkg:has_npc_pathway ?np_pathway .
          ?annotation enpkg:has_InChIkey2D ?ik_2d .
          ?feature enpkg:has_sirius_annotation ?annotation .
          ?feature enpkg:has_row_id ?row_id .
          ?annotation enpkg:has_cosmic_score ?cosmic .
          ?annotation enpkg:has_sirius_adduct ?adduct .
          ?feature_list enpkg:has_lcms_feature ?feature .
          ?lcms enpkg:has_lcms_feature_list ?feature_list .
          ?lcms enpkg:has_massive_doi ?massive_id .
          ?lcms rdf:type ?mode .
          ?sample enpkg:has_LCMS ?lcms .
          ?material enpkg:has_lab_process ?sample .
          ?material enpkg:has_wd_id ?species .
     }
```

### Supplementary Figures

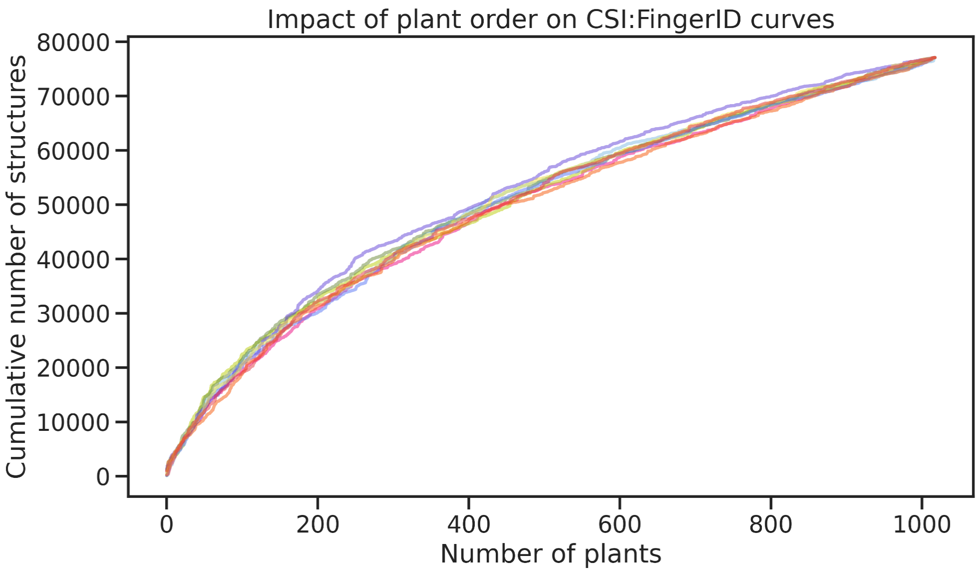


**Supplementary Figure 1**. **Shuffling the order of the plants does not significantly change the shape of the curve.** Because the curves in **Figure 3** all have similar shapes, the plant ordering differences will be similar for the other datasets. Each color represents a curve made in a different order. The curves look similar regardless of the methodology used (e.g., formulas from SIRIUS, MS2Mol predictions, etc). While minor variations in curve shape caused by shuffling the order may carry over to the projected curves in **Figure 5**, these differences will not meaningfully impact the range of the estimates, which is the primary focus of that figure.

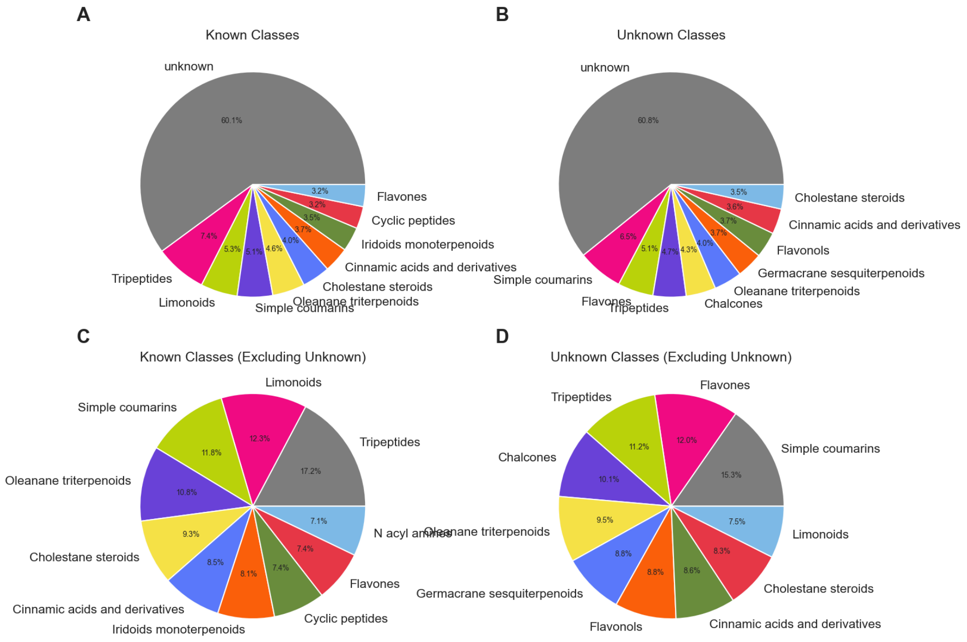


**Supplementary Figure 2. A) Distribution of NP classes for known (confidently annotated) structures. B) Distribution of NP classes for unknown (low confident annotations)**

**structures. C) Distribution of NP classes excluding unknowns among the confidently annotated structures. D) Distribution of NP classes excluding unknowns among the low confident annotated structures.**

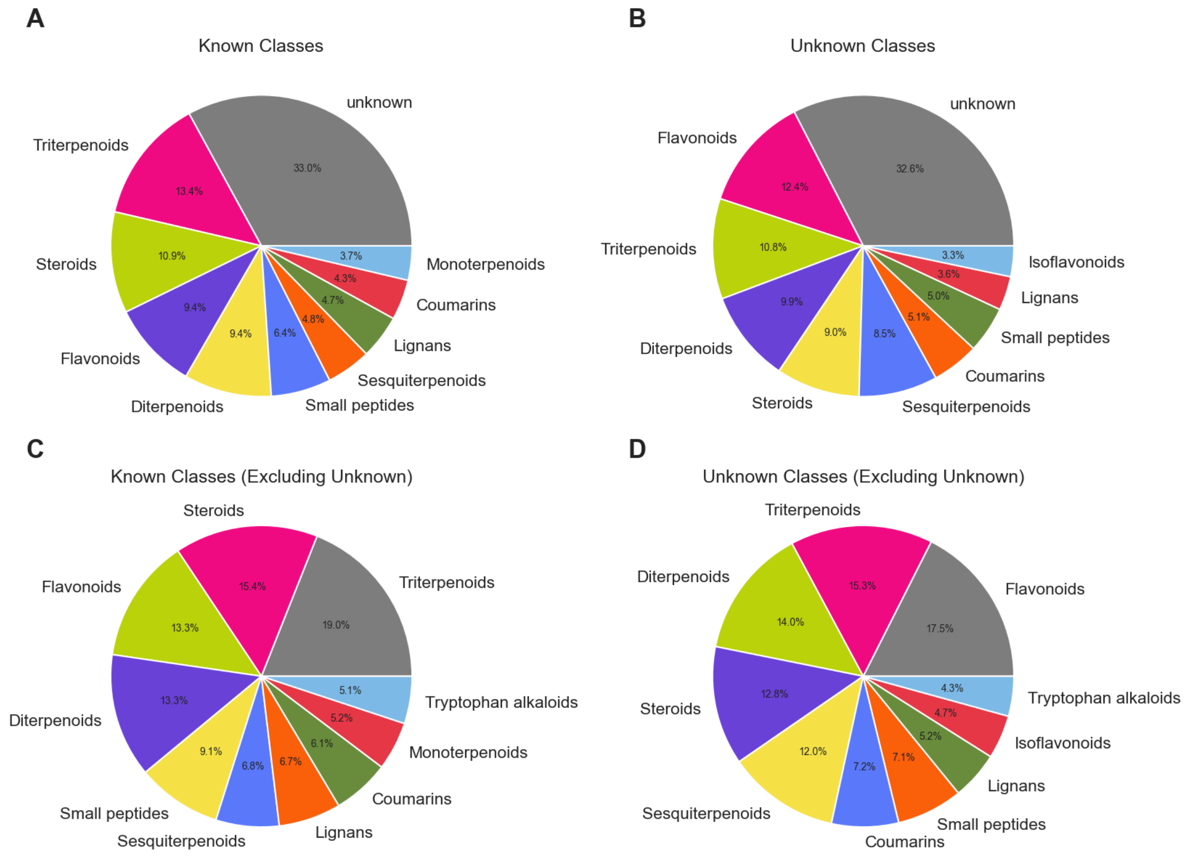


**Supplementary Figure 3. A) Distribution of NP super classes for known (confidently annotated) structures. B) Distribution of NP super classes for unknown (low confident annotations) structures. C) Distribution of NP super classes excluding unknowns among the confidently annotated structures. D) Distribution of NP super classes excluding unknowns among the low confident annotated structures.**

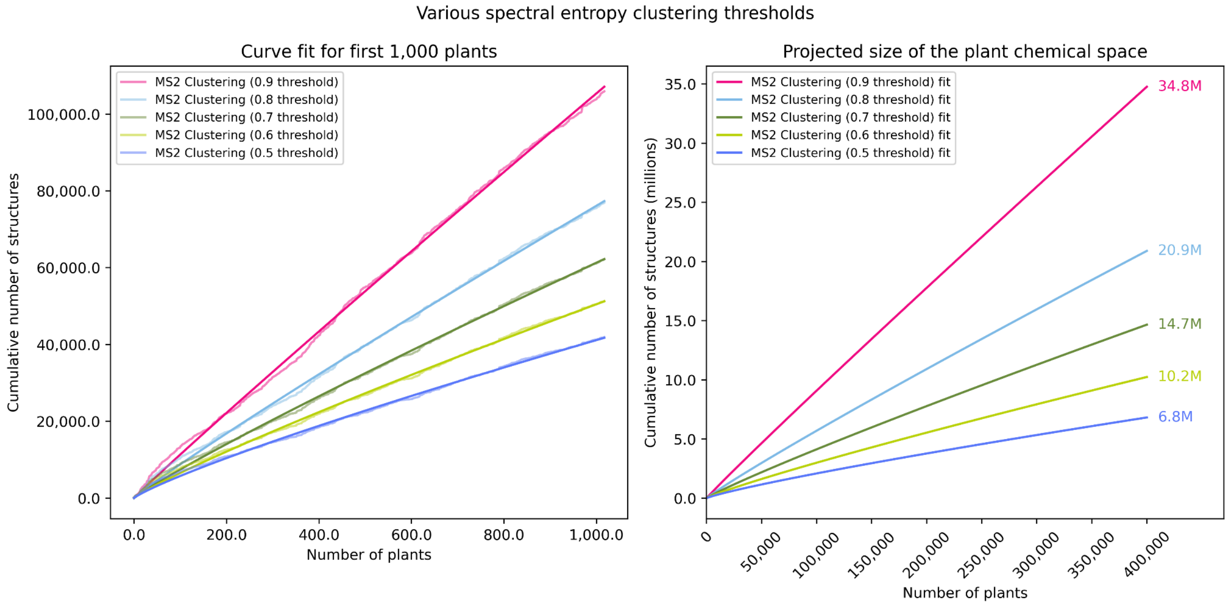


**Supplementary Figure 4.** Various spectral entropy thresholds were employed for MS2 clustering, resulting in projections for the number of metabolites in all plant space ranging from 6.8 million to 34.8 million. For the main results of our paper, we use the moderate estimates.

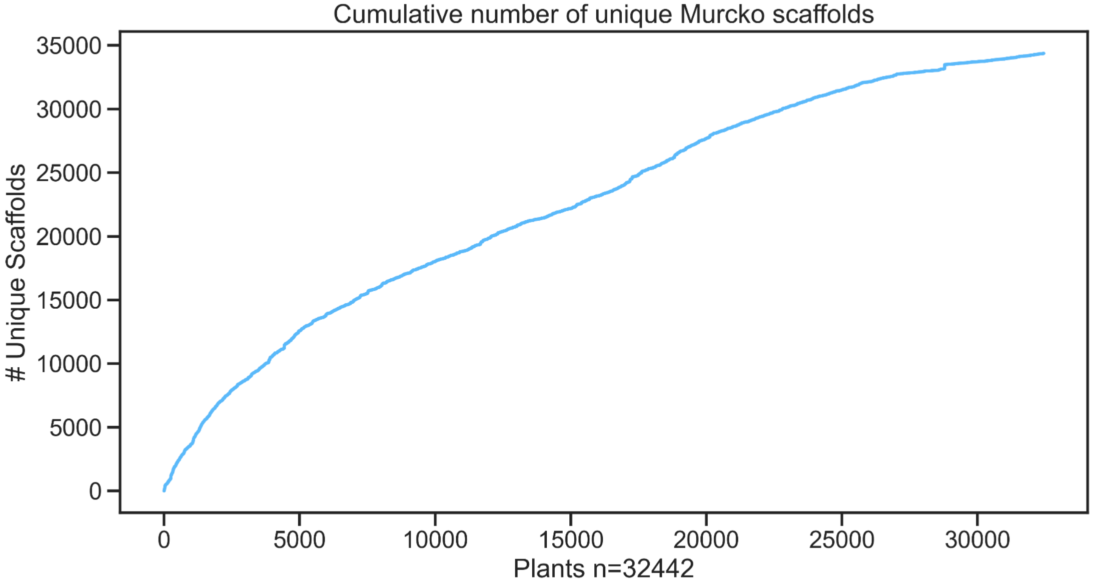


**Supplementary Figure 5. Cumulative curve of unique Murcko scaffolds in the literature dataset.** In total, there are 34,371 unique Murcko scaffolds in the literature datasets. Given that there are 124,086 unique structures, there is an average new Murcko scaffold for every 3.61 novel structures.

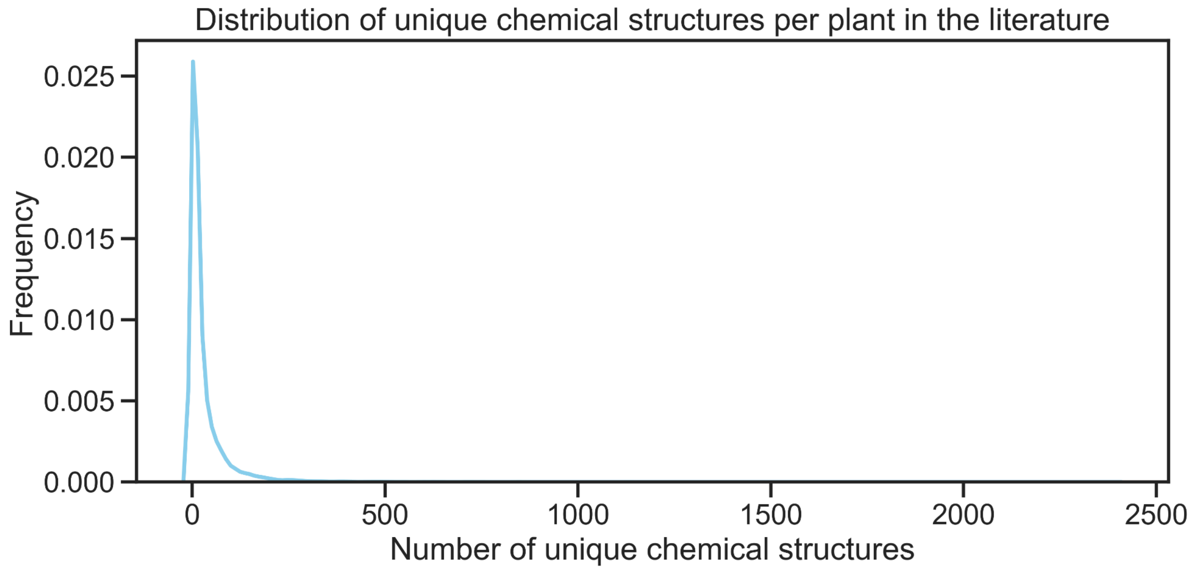


**Supplementary Figure 6 Distribution of unique chemical structures in the literature dataset.** The majority of the plants exclusively contain a few structures.

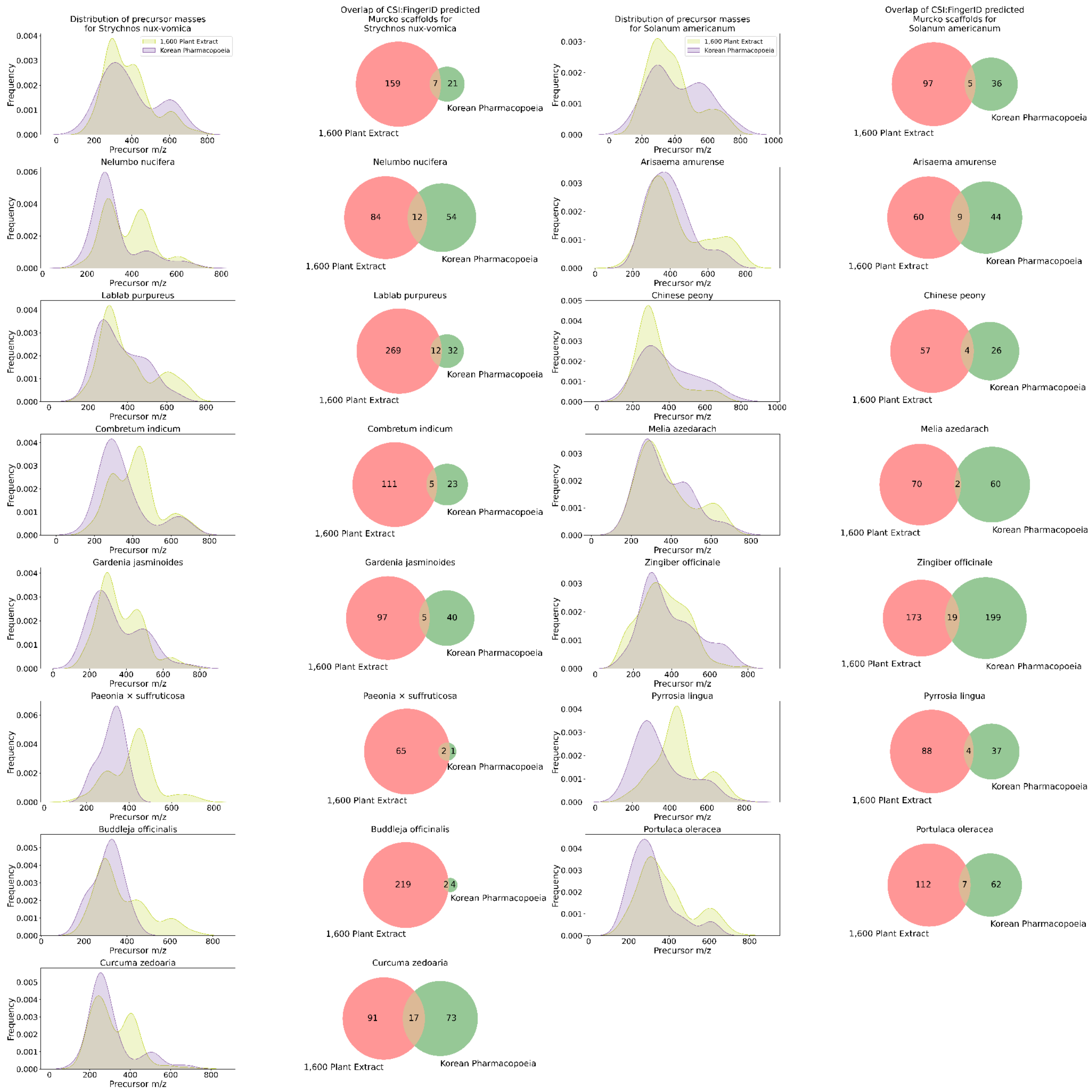


**Supplementary Figure 7. Distribution of precursor masses and overlap of Murcko scaffolds predicted with CSI:FingerID for the 15 plants present in both the Korean Pharmacopoeia dataset (Kang *et al.,* 2022) and the Pierre Fabre Research Institute Library (1,600 plant extracts) (Allard *et al.*, 2024) datasets.** We observe significant differences in the precursor masses distribution between the two datasets for the same plant and on the predicted Murcko scaffolds. Additionally, for these 15 plants present in both datasets, the extracts from the 1,600 plant extract dataset contain more features than those from the Korean Pharmacopoeia dataset. As discussed in the introduction, these differences are probably due to the different extraction methods used, among other factors. The high-quality version of this figure is present in the GitHub repository, as we could not upload it, retaining the original quality here.

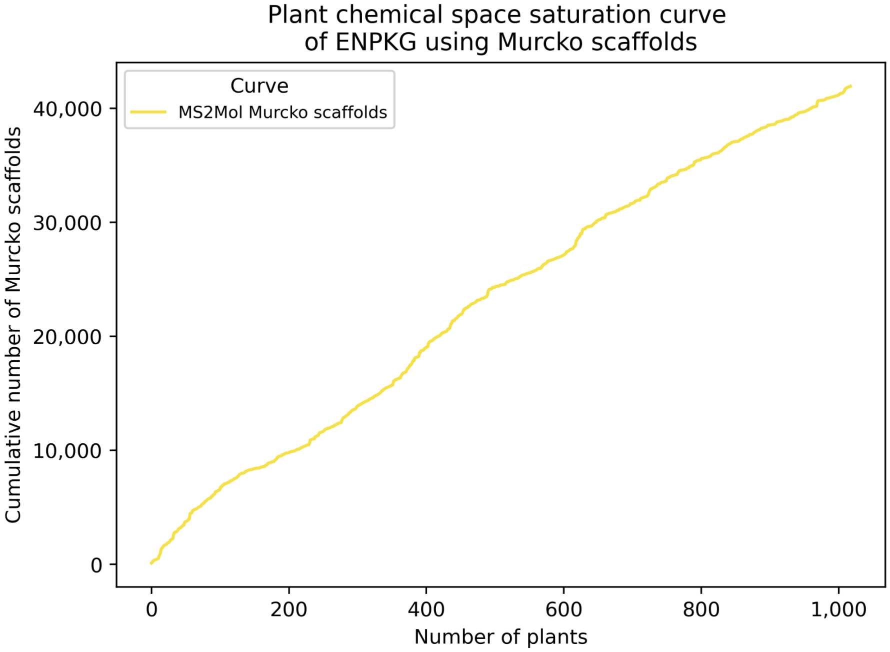


**Supplementary Figure 8. Plant chemical space saturation curve of ENPKG using Murcko scaffolds of the predicted structures from MS2Mol.**

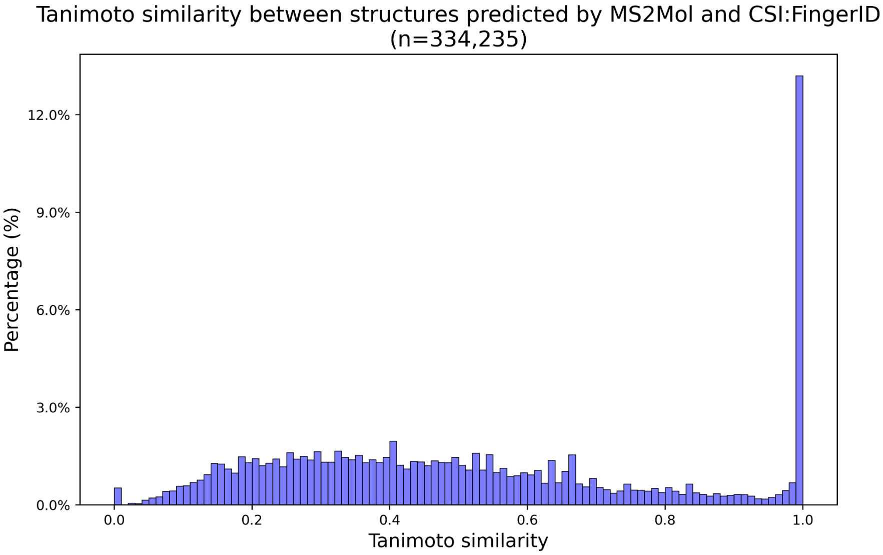


**Supplementary Figure 9. Distribution of Tanimoto similarity between the chemical structures predicted with MS2Mol and CSI:FingerID**. The Tanimoto similarity is calculated for each pair of predicted SMILES for a given MS2 using RDKit fingerprints with default parameters. The average Tanimoto coefficient is 0.51, the standard deviation is 0.28, the 25% percentile is 0.29, the 50% percentile is 0.45, and the 75% is 0.70.

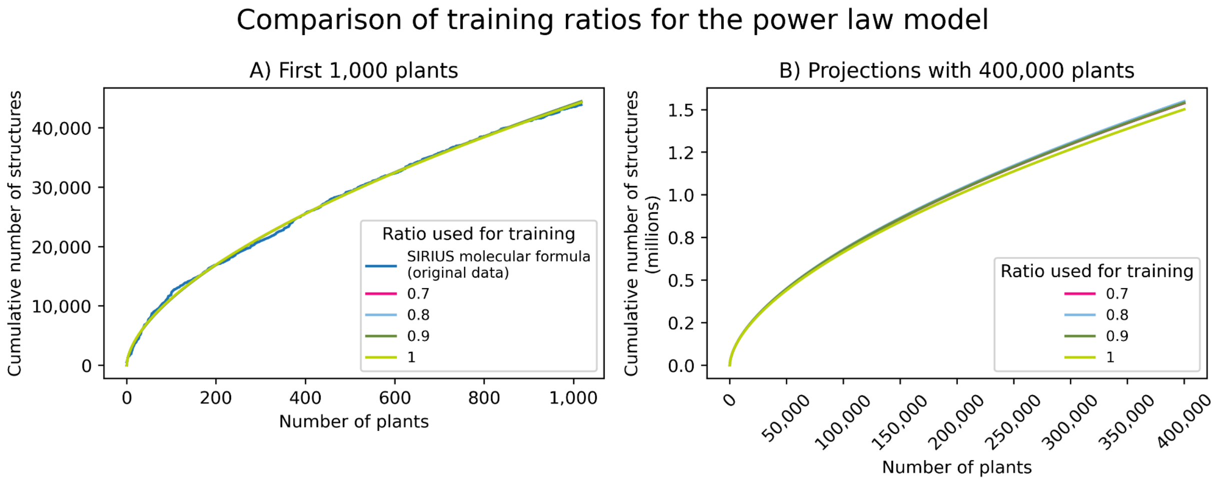


**Supplementary Figure 10. Comparison of training ratios for the power law model.** A) Curves fitted using different training ratios (0.7, 0.8, 0.9, 1) compared to the actual curve of the predicted formulas in the approximately 1,000 plants in the metabolomics dataset. B) Projection of the curves fitted in A) using different training ratios on the 400,000 species. Note that while the fitting ratio seems similar in the first 1,000 plants, these subtle differences translate into differences of several hundred thousand unique compounds when extrapolating to all 400,000 species.

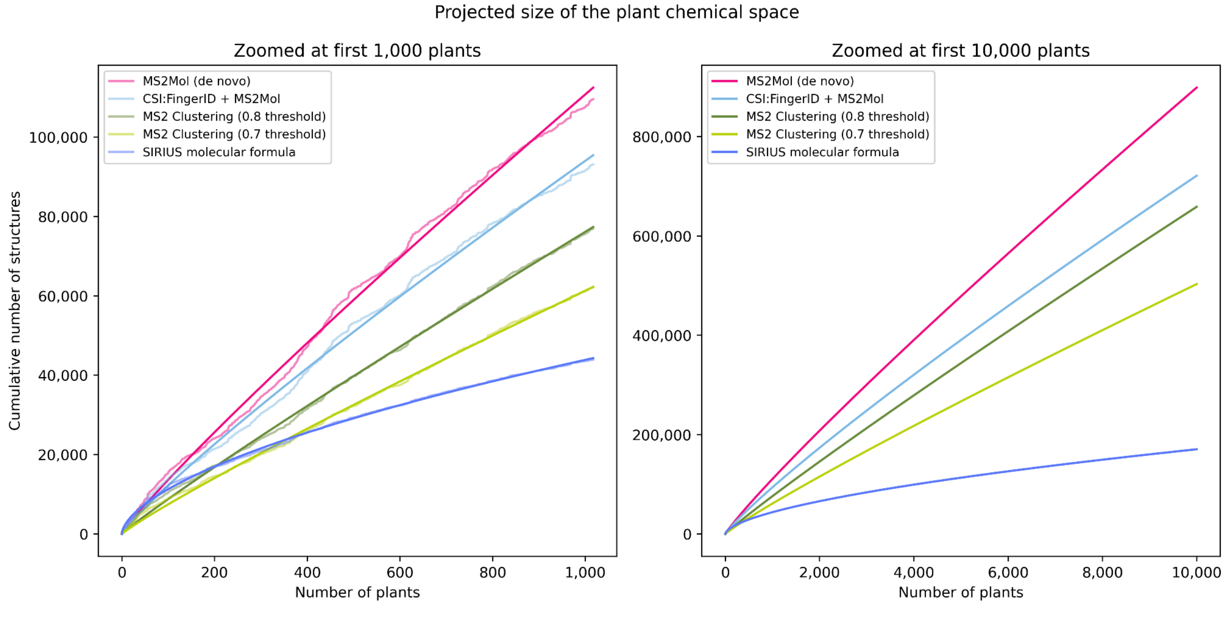


**Supplementary Figure 11. The projected size of the plant chemical space after fitting power law models on the different methodologies on the metabolomics dataset.** A) This plot is a zoomed-in version of **Figure 4** in the manuscript. We can see how the fitted curves for the projections fit the actual curves from the metabolomics datasets. The different assumptions/nuances across the methods explain the differences in the curve shape. B) When zoomed in to 10,000 plants, we can see that the CSI:FingerID + MS2Mol curve eventually crosses the MS2 Clustering (0.8 threshold) curve as the hybrid curve plateaus more significantly. This behavior is likely due to the influence of CSI:FingerID predictions, which are constrained by a reference library.

### Supplementary Tables

| Adduct | Number of features in ENPKG |
|---|---|
| [M+H]+ | 335,377 |
| [M-H]- | 259,955 |
| [M+NH4]+ | 152,603 |
| [M+Na]+ | 72,332 |
| [M+K]+ | 11504 |
| [M+FA-H]- | 4 |
| [M + O + H]+ | 3 |

**Supplementary Table 1. Distribution of each adduct predicted by SIRIUS for all 831,778 features in ENPKG.**

| Nomenclature | Unique species |
|---|---|
| NCBITax | 26,008 |
| ITIS | 1,019 |
| WFO | 5,415 |
| **Combined** | **32,442** |

**Supplementary Table 2. Number of unique plants in the combined literature dataset (COCONUT and LOTUS) after species harmonization.**

| Adducts |
| --- |
| [M+H]+ |
| [M+K]+ |
| [M+H3N+H]+ |
| [M-H2O+H]+ |
| [M-H4O2+H]+ |
| [M-H]- |
| [M+Cl]- |
| [M-H2O-H]- |
| [M+CH2O2-H]- |

**Supplementary Table 3. List of adducts used to calculate the potential precursor mz mass shifts that are later matched against the exact masses of reported compounds in the literature for a given plant.**

| Taxonomic level | Count |
| --- | --- |
| Order | 61 |
| Family | 181 |
| Subfamily | 113 |
| Genera | 672 |

**Supplementary Table 4. Number of taxonomic clades covered by the metabolomics datasets.** In total, the dataset covers 672 genera, which means, on average, there is approximately one new genus every six species. Similarly, there are a total of 181 families, which corresponds to approximately five species per family. The most common family is *Fabaceae*, which has 77 plants (7.5% of the total species). The most common genus is *Garcinia*, which has nine plants (0.88% of the total species).

### Supplementary Functions

$$y = ax^b$$

**Supplementary Function 1**. Power law model used to project plant metabolite count curves
